# A novel nonsense variant in *SUPT20H* gene associated with Rheumatoid Arthritis identified by Whole Exome Sequencing of multiplex families

**DOI:** 10.1101/430223

**Authors:** Maëva Veyssiere, Javier Perea, Laetitia Michou, Anne Boland, Christophe Caloustian, Robert Olaso, Robert Olaso, Jean-François Deleuze, François Cornelis, Elisabeth Petit-Teixeira, Valérie Chaudru

## Abstract

The triggering and development of Rheumatoid Arthritis (RA) is conditioned by environmental and genetic factors. Despite the identification of more than one hundred genetic variants associated with the disease, not all the cases can be explained. Here, we performed Whole Exome Sequencing in 9 multiplex families (N=30) to identify rare variants susceptible to play a role in the disease pathogenesis. We pre-selected 73 genes which carried rare variants with a complete segregation with RA in the studied families. Follow-up linkage and association analyses with pVAAST highlighted significant RA association of 24 genes (p-value < 0.05 after 10^6^ permutations) and pinpointed their most likely causal variant. We re-sequenced the 10 most significant likely causal variants (p-value ≤ 0.019 after 10^6^ permutations) in the extended pedigrees and 9 additional multiplex families (N=110). Only one SNV in *SUPT20H, c.73T>A (p.Lys25*)*, presented a complete segregation with RA in an extended pedigree with early-onset cases. In summary, we identified in this study a new variant associated with RA in *SUPT20H* gene. This gene belongs to several biological pathways like macro-autophagy and monocyte/macrophage differentiation, which contribute to RA pathogenesis. In addition, these results showed that analyzing rare variants using a family-based approach is a strategy that allows to identify RA risk loci, even with a small dataset.

**Author summary:** Rheumatoid arthritis (RA) is one of the most frequent auto-immune disease in the world. It causes joint swellings and pains which can lead to mobility impairment. To date, the scientific community has identified a hundred genes carrying variants predisposing to RA, in addition to the major gene HLA-DRB1. However, they do not explain all cases of RA. By examining nine families with multiple RA cases, we identified a new rare nonsense variant in *SUPT20H* gene, *c.73T>A (p.Lys25*)*. This finding is supported by the literature as the *SUPT20H* gene regulates several biological functions, such as macro-autophagy or monocyte/macrophage differentiation, that contribute to RA pathogenesis.

## Introduction

Rheumatoid Arthritis(RA) is one of the most frequent autoimmune disease, affecting 0.3 to 1% of the worldwide population. Since the discovery of HLA locus as a risk factor for autoimmune diseases [1] and specifically HLA-DRB1 for RA, more than 100 RA genetic factors were identified by Genome Wide Association Studies (GWASs) [2,3]. However, the effect of these genetic risk factors is too weak to explain the entire RA genetic component. Indeed, the heritability attributed to HLA-DRB1 shared-epitope (SE) alleles was estimated between 11% [4] and 37% [5]. While GWASs loci identified outside the HLA locus only explain an additional five percent of RA heritability [6].

Several hypotheses have been proposed to explain the unknown part of this complex disease genetic component. One of these hypotheses relies on the fact that rare variants, which are poorly detected by GWASs, contribute to the risk of complex diseases. During the last decade, the development of Next Generation Sequencing has facilitated the detection of such variants. Hence, several studies used exome sequencing to identify rare to low frequency variants and evaluate their contribution to RA risk. For this purpose, two studies used a candidate gene approach based on exome sequencing [7,8]. A first study, based on a population of European ancestry, showed an aggregation of non-synonymous rare variants contributing to RA risk into IL2RA and IL2RB loci [7]. Another one relying on Whole Exome Sequencing (WES), which studied Korean RA cases and healthy controls, allowed the identification of a weak association of rare missense variants in 17 different genes [8]. In this latter group, the VSTM1 gene gave rise to an over-expression of its mRNA product in RA cases [9]. However, both studies restricted their analysis to a limited number of candidate genes. Thus, they may have missed variations contributing to RA risk present outside the loci of those candidates. Other studies [10,11] succeeded at identifying new RA risk loci by analyzing rare coding variants extracted after Whole Exome Sequencing. Thus, an association between RA and *PLB1* gene [11] and several other genes involved in the production of reactive oxygen species (ROS) such as NDUFA7 and SCO1 [10] has been observed using respectively a familial-based strategy and a case-control analysis. Both studies, conducted in non-European populations, support the strategy of sequencing the whole exome of RA cases to identify new RA candidate variants.

In our study, we aimed at identifying new loci associated with RA in the French population by focusing our research on rare coding variants. For this purpose we sequenced RA cases and healthy relatives from 9 multiplex families which carried HLA-DRB1 risk alleles. However, even in the cases who carried the SE alleles, we can observe some clinical and genetic heterogeneity. Hence, we should be able to identify both genetic factors modulating the effect of HLA-DRB1 SE and acting independently from HLA loci.

## Results

### Overview

In this study, we sequenced the whole exome of 19 RA cases and 11 healthy individuals belonging to 9 multiplex families (discovery set). We applied a three steps strategy (Fig 1) to prioritize the sequenced loci and identify RA risk variants. First, we selected genes carrying rare variants which were family specific and with a high damaging predicted effect. Then, we assessed the potential combined effect of rare and low-frequency variants in these genes on RA risk using pVAAST [12]. Finally, for the 10 genes with the strongest genetic association, we re-sequenced the leading effect variant in the validation set. This final step allowed us to validate their family specific co-segregation with RA.

**Fig 1:**
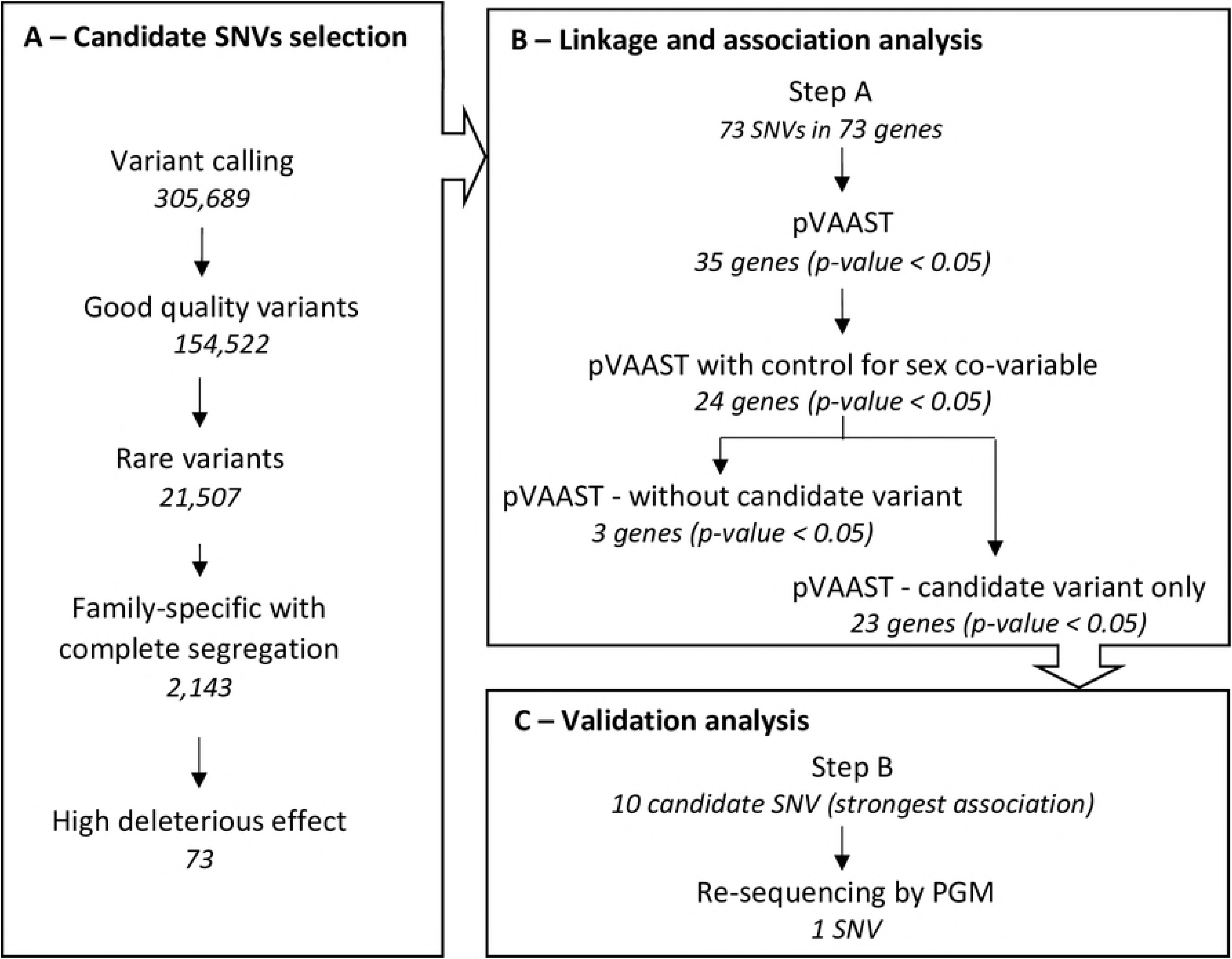
Description of the study design. This schema of the study presents the 3 main steps of the analysis and, the resulting number of variants selected through each sub-analysis. (A) The selection of candidate variants was performed on the discovery set and resulted in the identification of 73 candidate SNVs (B) The association analysis was applied on the discovery set and 45 controls extracted from the 1000 genomes project. The 24 genes significantly associated with RA were re-tested after removing the candidate variant identified in step A and, the candidate variant was tested alone (C) The SNVs with the strongest RA association were re-sequenced in the extended families of the discovery, plus 9 new multiplex families.

### Description of the discovery set

The male: female ratio was 6:13 in the discovery set selected for WES (described in Table 1), which is similar to the ratio observed in RA affected populations [13,14]. All the RA-affected individuals carried at least 1 shared epitope allele of HLA-DRB1 (18 of 19 had 2 SE alleles). Fifteen affected participants (79%) were positive for Anti-Citrullinated Peptide Antibodies (ACPA) and Rheumatoid Factor (RF), one (5%) was positive for the RF alone and two (11%) were negative for both. These two seronegative cases belonged to the same family (referred as family 5). The affected members of families 3, 4 and 9 were diagnosed with early RA (the mean age at diagnosis was of 31, 36 and 26 years old, respectively).

**Table 1:**
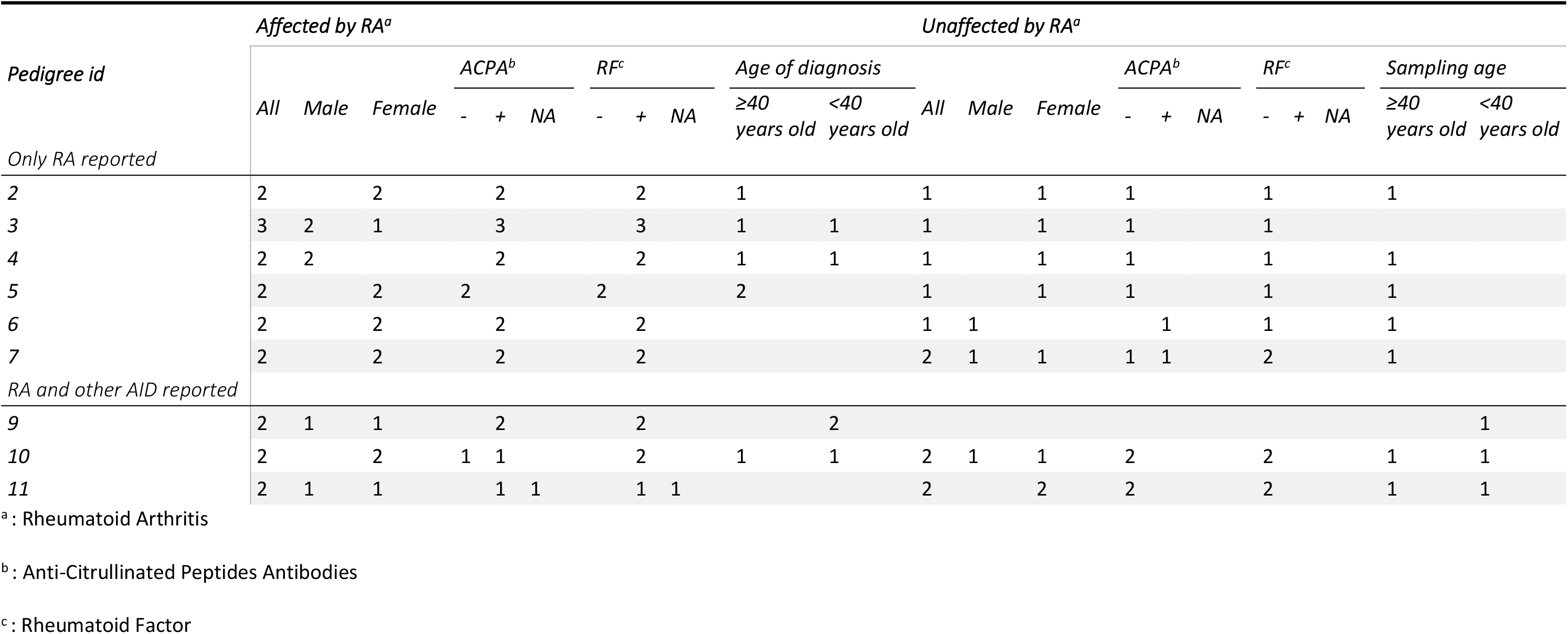
Description of the discovery set.

To check whether the 45 selected controls would not inflate the type I error during the burden test, we performed a PCA based on these samples and the discovery set. The genetic variability (<3.6% of the total variability) observed in the PCA (Fig 2) was due to the variability between the selected families and not between families and controls.

**Fig 2:**
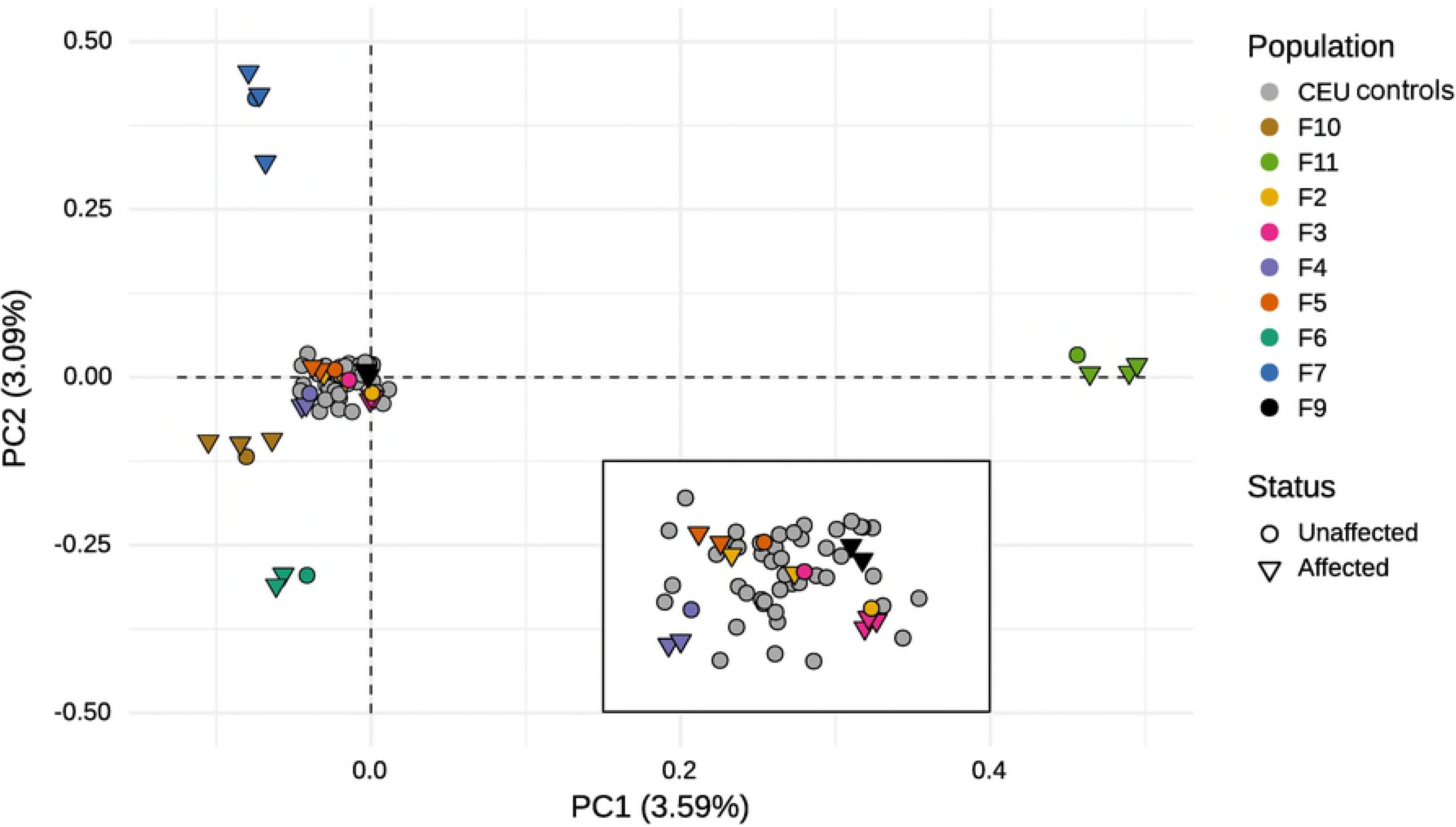
Principal component analysis (PCA) results of samples processed for pVAAST analysis. The PCA is based on the 30 samples from the discovery set and 45 CEU controls extracted from the 1000 genomes databases. This plot represents the 2 first principal components which respectively account for 3.59% and 3.09% of the genetic variability. The color code represents the population source: each family sequenced by WES has its own color described in the legend, the CEU controls are in grey. The shape of each dot represents the RA status of the represented individual.

### Whole exome sequencing and RA risk candidate variants detection

More than 91% of the targeted regions were sequenced with a coverage ≥ 30X. We extracted from these sequences 154,644 high-quality variants, including 64,747 (42%) in exons. We observed a mean of 26,679 exonic variants per sample and a SNV transition/transversion ratio equal to 3.01 which is consistent with previous studies [15].

Under the hypothesis that rare genetic variants linked to RA would segregate with the affected phenotype within the multiplex families, we selected in the pool of 21,507 rare variants (MAF < 1%), 2143 family-specific variants shared by all the RA-affected relatives (but absent from unaffected members). Knowing that rare variant with high predicted biological effect may contribute to the genetic predisposition of common disease, we extracted, from the 2143 variants, 73 SNVs with a high deleterious predicted effect on protein. We based our evaluation of this impact on SNPeff (“HIGH” or “MODERATE”) and CADD phred score (≥30). All these 73 rare family-specific variants were heterozygous in affected carriers. They were detected in 73 different genes, not previously associated with RA.

### Evaluation of candidate variant association with RA

We reduced the set of candidate loci to 35 genes (48% of the genes tested with pVAAST) which were significantly associated with RA (pVAAST_p-value_ < 0.048 after 10^6^ permutations – Table 2). Considering the difference in RA prevalence rate between men and women, we performed pVAAST test while controlling for the variable “sex”. A total of 24 genes still had a significant association with RA (p-value < 0.05 – see Table 2).

**Table 2:**
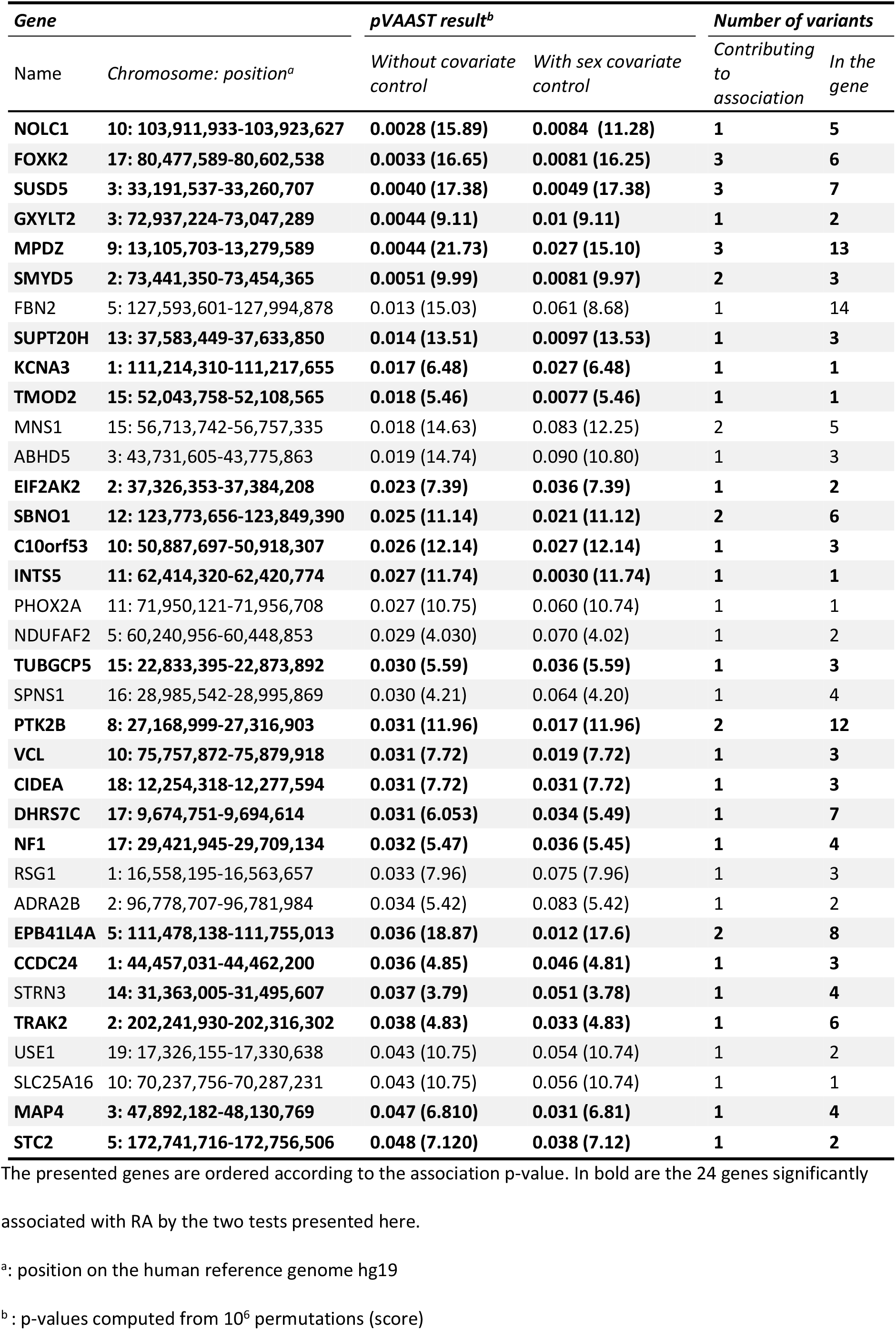
Genes showing association signal in pVAAST analysis.

We categorized the 24 genes into two groups by looking at the association scores of their variants (Table 3). Seven genes (group 1) carried several variants, including our best rare candidate SNV, which contributed to the score of the gene (SNV_score_ >0). In the other seventeen genes (group 2), only the family-specific variant participated to the gene score. In the 24 genes, the rare family-specific variants had the highest score. To validate the leading effect of the candidate variants, we performed again the burden test by including first, only the candidate SNVs, and then, all variants except these candidates (results in Table 3). All the 24 candidate SNVs, except the SUSD5:c.526C>T (p-value = 0.23) were significantly associated with RA (p-value ≤ 0.048). Concerning the test without the candidate SNV, three genes in the first group (22%), *SUSD5, MPDZ and FOXK2*, were still significantly associated with RA (p-value < 0.05). So, several rare variants in our dataset contributed to the association of these genes with RA. However, none of the genes in the second group were associated with RA after excluding the candidate variant. This observation confirmed that the observed RA-association with these genes was in fact due to the family-specific variant.

**Table 3:** Association analysis of candidate SNVs in 24 candidate genes.

| *Gene* | *Chromosome: position<sup>a</sup>* | *Category<sup>b</sup>* | *Candidate variant (CV)* | *pVAAST<sup>c</sup>* |  |
| --- | --- | --- | --- | --- | --- |
|  |  |  |  | *CV only* | *Without CV* |
| SUPT20H | 13: 37,583,449-37,633,850 | 2 | c.73T>A | 0.0029 (13.53) | 1 (0) |
| NOLC1 | 10: 103,911,933-103,923,627 | 2 | c.331C>T | 0.0041 (11.28) | 0.20 (2.61) |
| GXYLT2 | 3: 72,937,224-73,047,289 | 2 | c.358C>T | 0.0046 (9.11) | 1 (0) |
| TUBGCP5 | 15: 22,833,395-22,873,892 | 2 | c.2327G>A | 0.015 (5.59) | 1 (0) |
| MAP4 | 3: 47,892,182-48,130,769 | 2 | c.2987C>T | 0.016 (6.81) | 1 (0) |
| EIF2AK2 | 2: 37,326,353-37,384,208 | 2 | c.1078G>A | 0.016 (7.39) | 1 (0) |
| PTK2B | 8: 27,168,999-27,316,903 | 1 | c.29G>A | 0.016 (4.83) | 0.26 (5.13) |
| DHRS7C | 17: 9,674,751-9,694,614 | 2 | c.605T>C | 0.017 (5.37) | 0.38 (0.71) |
| TMOD2 | 15: 52,043,758-52,108,565 | 2 | c.263C>T | 0.018 (5.46) | 1 (0) |
| STC2 | 5: 172,741,716-172,756,506 | 2 | c.131G>T | 0.019 (7.12) | 1 (0) |
| NF1 | 17: 29,421,945-29,709,134 | 2 | c.7964C>T | 0.021 (5.45) | 1 (0) |
| SBNO1 | 12: 123,773,656-123,849,390 | 1 | c.2170G>T | 0.022 (8.61) | 0.48 (0.51) |
| TRAK2 | 2: 202,241,930-202,316,302 | 2 | c.1526G>A | 0.023 (4.83) | 1 (0) |
| INTS5 | 11: 62,414,320-62,420,774 | 2 | c.1663C>T | 0.023 (11.74) | 1 (0) |
| VCL | 10: 75,757,872-75,879,918 | 2 | c.1558C>T | 0.024 (7.72) | 1 (0) |
| C10orf53 | 10: 50,887,697-50,918,307 | 2 | c.195C>A | 0.025 (12.14) | 1 (0) |
| EPB41L4A | 5: 111,478,138-111,755,013 | 1 | c.1828C>T | 0.026 (12.14) | 0.32 (5.19) |
| KCNA3 | 1: 111,214,310-111,217,655 | 2 | c.640C>T | 0.027 (6.48) | 1 (0) |
| CIDEA | 18: 12,254,318-12,277,594 | 2 | c.112C>T | 0.03 (7.72) | 1 (0) |
| SMYD5 | 2: 73,441,350-73,454,365 | 1 | c.833G>A | 0.031 (5.59) | 0.12 (2.38) |
| CCDC24 | 1: 44,457,031-44,462,200 | 2 | c.848G>A | 0.037 (4.81) | 1 (0) |
| FOXK2 | 17: 80,477,589-80,602,538 | 1 | c.1321C>T | 0.044 (6.32) | 0.044 (8.33) |
| MPDZ | 9: 13,105,703-13,279,589 | 1 | c.6022A>T | 0.048 (2.76) | 0.021 (16.96) |
| SUSD5 | 3: 33,191,537-33,260,707 | 1 | c.526C>T | 0.23 (3.94) | 0.013 (11.45) |

The genes are sorted according to the association p-value of the candidate variant.
<sup>a</sup>: position on the human reference genome hg19
<sup>b</sup>: The two categories correspond to:
1 – genes containing multiple rare variants contributing to pVAAST score
2 – genes containing one rare variant leading the association
<sup>c</sup>: p-values computed from 10<sup>6</sup> permutations (score)

### Validation in extended pedigrees

We selected the top 10 RA candidate variants, with the strongest association with RA, for re-sequencing to confirm their family specificity and their co-segregation with RA (Table 4). We analyzed further 9 out of 10 variants for which we were able to produce sequences and, we validated the familial specificity for all of them. Five variants (in *SUPT20H, NOLC1, GXYLT2, MAP4 and STC2* genes) were shared by all affected members in the families who carried these variants. However, only one candidate variant showed a complete segregation with RA. This variant is a heterozygous non-sense SNV, *c.73T>A (p.Lys25*)*, located in *SUPT20H* gene which introduces a premature codon stop at the beginning of the gene (in the fourth exon) (Fig 3). This variant, not reported to date, has been deposited in ClinVar database prior to publication.

**Table 4:** Top 10 RA associated SNVs and their re-sequencing results.

| *Gene* | *Chromosome* | *Position (bp)* | *Ref/Alt* | *rs number<sup>a</sup>* | *Family* | *#RA cases* | *#Unaffected* |
| --- | --- | --- | --- | --- | --- | --- | --- |
| SUPT20H | 13 | 37622040 | T/A | - | 3 | 4/4 | 0/5 |
| NOLC1 | 10 | 103917202 | C/T | - | 3 | 4/4 | 2/5 |
| GXYLT2 | 3 | 72957600 | C/T | rs375437631 | 3 | 4/4 | 1/5 |
| TUBGCP5 | 15 | 22864369 | G/A | rs138440581 | 2 | 2/3 | 1/3 |
| MAP4 | 3 | 47908815 | G/A | rs150907099 | 5 | 5/5 | 2/3 |
| EIF2AK2 | 2 | 37347149 | C/T | - | 5 | 4/5 | 1/3 |
| PTK2B | 8 | 27255130 | G/A | rs111853465 | 4 | 2/5 | 4/6 |
| DHRS7C | 17 | 9676206 | A/G | rs115900148 | 4 | NA | NA |
| TMOD2 | 15 | 52060595 | C/T | - | 4 | 4/5 | 3/6 |
| STC2 | 5 | 172755066 | C/A | rs148833559 | 2 | 3/3 | 1/3 |

<sup>a</sup>: rs number from dbsnp138

**Fig 3:**
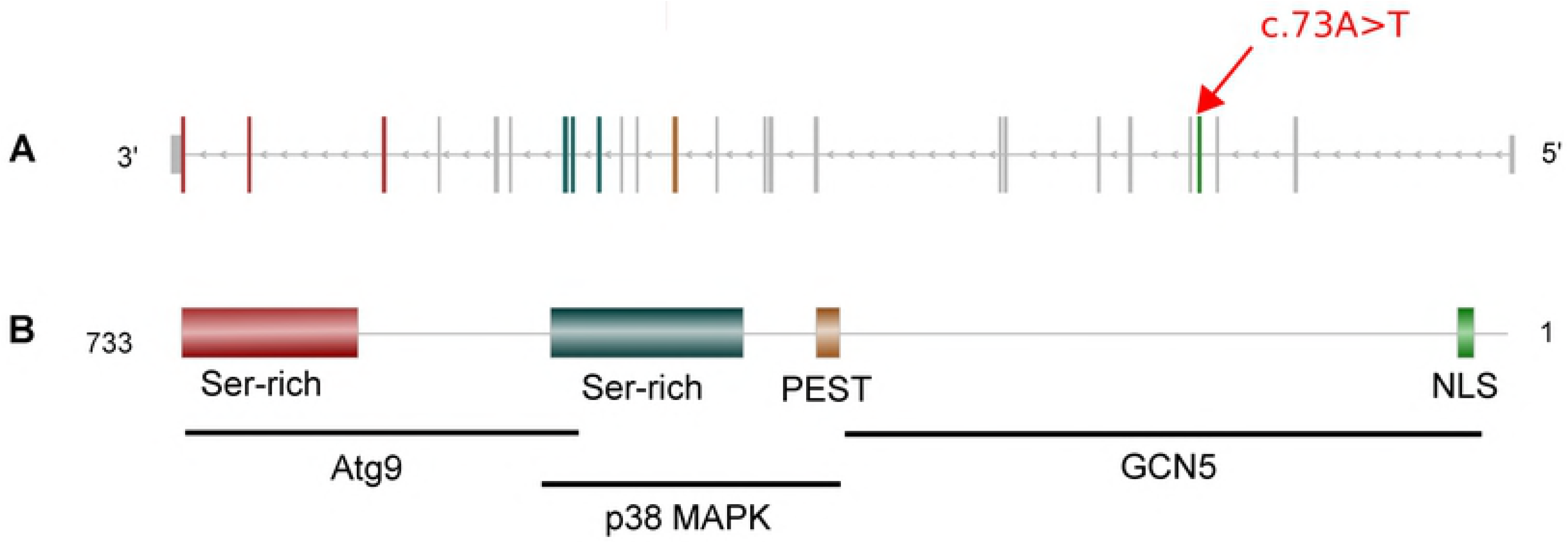
Gene SUPT20H and its product: p38IP. Abbreviations: NLS: Nuclear Localization signal, PEST: Proline (P) Glutamic acid (E) Serine (S) Threonine. (A) Gene model of SUPT20H. The red arrow indicates the exon including the nonsense variant identified by WES. The exons color code refers to the protein domain it encodes for. (B) On top, p38IP protein model of 733 amino-acid and on the bottom, the positions of p38IP interactors binding sites.

### Genotyping of SNV c.73T>A (p.Lys25*) in family 3 and in trios

The SNV *c.73T>A (p.Lys25*)* carried by *SUPT20H* was genotyped using a custom assay and digital PCR. First, genotyping results in family 3 validated the presence of the rare A allele in affected members of the family and, its absence in unaffected members (Fig 4). Second, we investigated 188 RA patients (characteristics described in supplementary Table S1) and 362 healthy parents from trio families using one affected member of family 3 as a positive control. No one of these samples showed the rare allele of this SNV.

**Fig 4:**
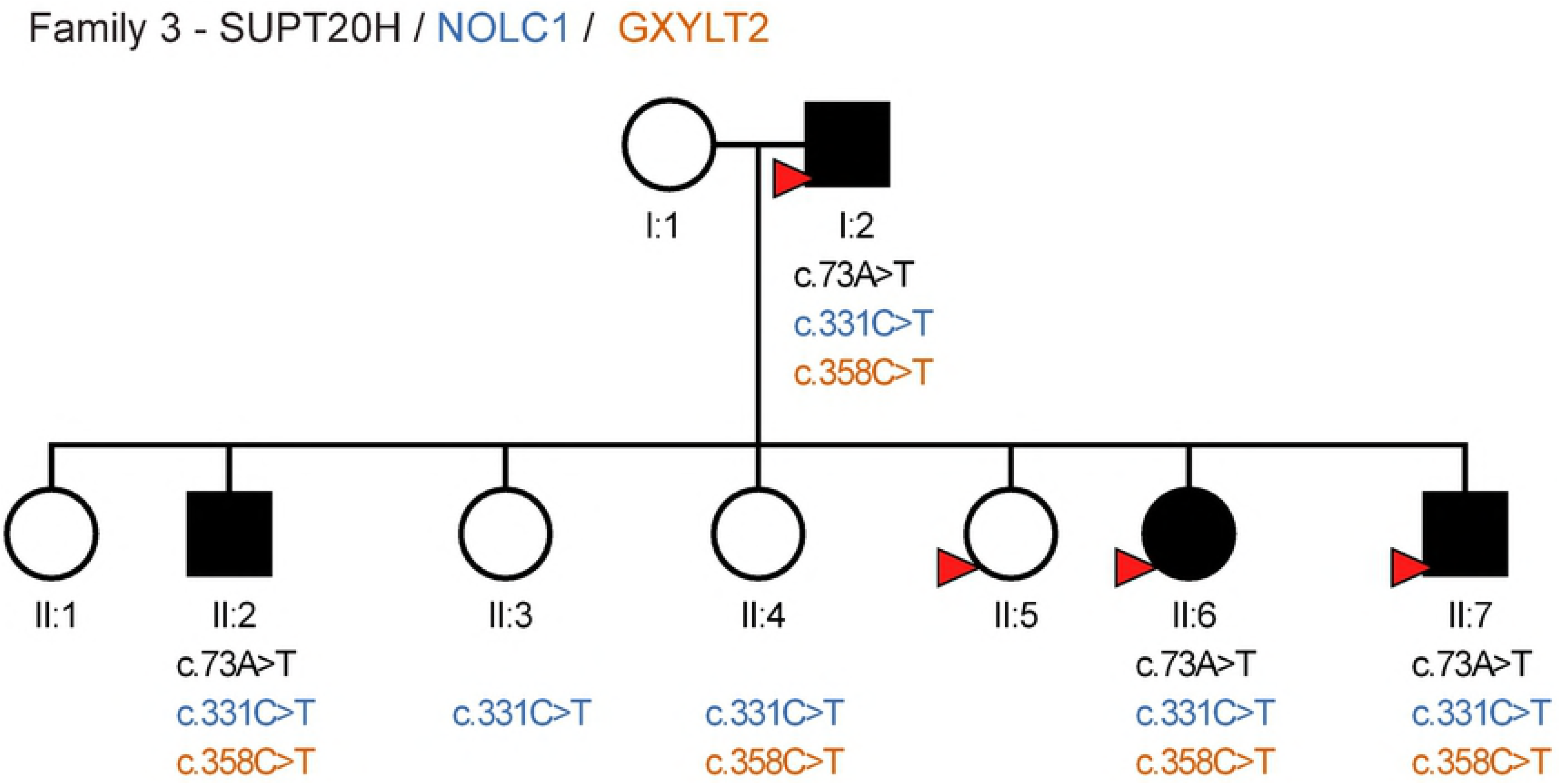
Representation of the pedigree 3 and the variants identified in his members. All the individuals represented here were part of the validation set but only the ones marked with a red arrow were part of the discovery set. Each of them are represented by a circle (if female) or a square (if male). Their filling color corresponds to their status: black if affected by RA or white if unaffected by the disease. The presence of a variant is represented by its HGVS id. The absence of this id indicate a homozygote genotype for the reference allele.

## Discussion

In this study, we identified a novel SNV associated with RA, SUPT20H: *c.73T>A (p.Lys25*)*, by performing WES in 9 multiplex RA pedigrees associated with HLA-DRB1-SE risk alleles. This nonsense variant had a complete penetrance in a family with rather young age at RA onset (mean_onset-age_ = 31 years old). In addition, we showed with the RVAT pVAAST, that this variant was significantly associated with RA (p-value = 2.9*10^-2^).

The *SUPT20H* gene, which has not been reported yet as a RA risk gene, is a member of the SAGA complex (Spt-Ada-Gcn5 acetyltransferase) gene family. It encodes for the p38 interacting protein (p38IP) which is predicted to contain a Nuclear Localization Signal (NLS), a PEST domain (Proline Glutamic acid Serine Threonine) and 2 serine rich domains [16]. The SNV identified in this study occurring in the NLS domain, the truncated protein that could result from this variation would not have any functional domain.

Previous studies reported p38IP interactions with 3 different proteins [16–18]. Indeed, the protein p38IP was shown to bind to and stabilize the protein GCN5 [17,19], member of the SAGA complex, and thus stabilize the complex itself [20]. In addition, *in vitro* analysis by Nagy and his colleagues [20] showed that this p38-GNC5 SAGA complex binds to the promotor of Endoplasmic Reticulum stress-induced genes, such as GRP78, enhancing their transcription. Interestingly, previous studies have noted the role of the GRP78 gene in the pathogenesis of RA: the expression of this gene is increased in RA fibroblast-like synoviocytes (FLS) and it promotes their proliferation [21,22]. The protein p38IP, as indicated by its name, interacts also with p38 MAPK, a key protein for the development of RA [23]. And it has previously been reported that this interaction inhibits monocyte/macrophage differentiation [19]. Macrophages, present in a high concentration in inflamed synovial membrane and cartilage/pannus junction, contributes to the maintenance of the inflammation in RA patients [24,25]. Finally p38IP interacts with the protein Atg9 and is required for its translocation from trans-Golgi system to endosome during starvation induced macro-autophagy [16,26,27]. The protein p38 MAPK, which has higher affinity for p38IP than Atg9 inhibits this interaction and thus inhibits the autophagy process. This implication of p38IP in autophagy process is another argument that highlights the interest of studying the role of p38IP in the development of RA. Indeed, different studies reviewed by Dai and Hu demonstrated the role of autophagy in RA [28].

In addition to SUPT20H gene, we identified 23 genes significantly associated with RA (p-value < 0.05). We evaluated their association with the test implemented in pVAAST. This software combines a rare variant association test (RVAT) with a linkage score, thus it takes into account for the familial structure of our data. It allowed us to provide all the variants called in our samples without any *a priori* filtering based on biological knowledge, avoiding loss of information. Indeed, the RVAT incorporates a score evaluating for each variant the amino acid substitution impact and phylogenetic conservation information, allowing it to be robust in the presence of neutral and common variants [12]. Indeed, depending on the chosen test, variants with opposite effects can lead to a decreasing power for the association test [29].

Among these 24 genes, we observed a variable number of variants contributing to the association signal: 7 genes with more than one contributing variant and 17 with only one. In the first group of genes, the potentially pathogenic variants covered the rare to low frequency spectrum. Three of these genes (named *FOXK2, SUSD5* and *MPDZ*), remained significantly associated with the disease without the candidate variants (p-value < 0.05). The fact that the signal for the other genes was not significant anymore, when tested without the candidate SNV, could be explained by the lower number of contributing variants identified in these genes that reduce the RVAT power compared to genes with more causal variants [30] (Table 2)In the second group, 3 genes (TMOD2, *INTS5* and *KCNA3*) contained only one variant. So, as expected, the score of the association test with all identified variants (familial data and control data) and with the candidate variant were identical (see Table 1).

We can highlight two limitations in this association analysis. First, we added as controls the variants of 45 individuals extracted from the 1000 genomes project database. And, although we chose individuals from the same population as our pedigrees (European) and sequenced on the same platform (Illumina), two aspects could have introduced bias: the exome capturing kit and the variant calling workflow. The first aspect can introduce variants specific to one of the dataset because not targeted by the kit used for the other one. To partially overcome this issue, we filtered out variants from the 1000 genomes dataset present outside the boundaries of our targeted exomes. But, due to the wide range of capturing kits used to produce the data presents in the 1000 genomes database, we did not filter the variants according to the regions targeted by the 1000 genomes project. For the second aspect, it has previously been shown that despite a few variant specific to a variant calling software, the concordance between the called variants is high (92% between HaplotypeCaller [31], Samtools [32] and FreeBayes [33])[34]. In addition, the PCA, performed on all the individuals included in the association test (Figure 2), shows that the observed genetic variability is mostly related to the intra-family specificity, not to the choice of the 45 CEU controls, comforting our choice of controls. Second, individuals of the discovery set have different clinical characteristics (such as sex, age at diagnostic, and ACPA status). These differences, if not controlled, could have introduced bias in the association analysis. Since, the sex was the sole information known for both discovery set and CEU controls, we were only able to control for this covariate.

Finally, we previously suggested that rare variations present in the genome of RA cases in addition to HLA-DRB1 SE could modulate the effect of this latter, leading to RA cases with different clinical characteristics. To test this hypothesis, we could investigate the interactions between HLA-DRB1 and new RA risk genes, such as SUPT20H. For this analysis, we would need to perform a study including both individuals carrying and not carrying HLA-DRB1 risk alleles. Further work is also needed to identify new RA risk genes that may have been missed in this study because we used stringent criteria to select the candidate SNVs. Indeed, we removed variants: with incomplete penetrance, segregating in several families and not shared by all affected relatives of a given family.

In conclusion, we identified a new rare nonsense SNV, SUPT20H: *c.73T>A (p.Lys25*)*, associated with RA, by combining linkage and association analysis with pVAAST. Neither the gene nor the variant were previously associated with the disease. But the review of the literature about SUPT20H gene and its product, p38IP, supports the idea that this gene is involved in the pathogenesis of RA. Further work, with *in vitro* functional studies, needs to be done to evaluate the pathogenic effect of this new SNV and to validate its role in the different processes previously described in the context of RA. In addition, further studies need to be done to validate the other RA risk loci identified in this study, in particular *FOXK2, MPDZ* and *SUSD5* genes in which we observed an aggregation of rare variants.

## Methods

### Participants

We studied 16 French RA multiplex families with at least 4 affected individuals per family carrying one or two HLA-DRB1 shared epitope (SE) allele. Half of the families had only RA cases, whereas the other half had RA and/or other Autoimmune Diseases (AID: Lupus Erythematosus, Vitiligo, Sjögren syndrome, Hashimoto’s thyroiditis) cases. Among the 110 recruited individuals within these families, 33 (30%) were reported as only affected by RA, 17 (15%) by RA and another AID, 15 (14%) by an AID different from RA and 45 (41%) were reported as unaffected. As discovery set, we selected 30 individuals belonging to 9 of the 16 previously described pedigrees, with a sufficient quantity of DNA for WES. This sample set consisted in 19 individuals with RA and 11 relatives not affected by RA (at least one per pedigree). All the remaining samples were included in the validation set.

Our study was approved by ethics committees of Hôpital Bicêtre and Hôpital Saint Louis (Paris, France; CPPRB 94–40). Each individual provided a written informed consent for the participation in the study.

For statistical analysis with the Rare Variant Association Test (RVAT), we added 45 healthy controls extracted from the 1000 genomes project database. We chose CEU individuals (Utah residents with Northern and Western European ancestry) for which whole exome sequences were generated by the use of an Illumina platform.

### Whole exome sequencing and variant calling

The exons were captured in the discovery set with Agilent SureSelect Human All Exon kit (V5) which target the exome of more than 20,000 genes. Then, we sequenced those regions on an Illumina HiSeq2000 platform and mapped the reads to the human reference genome hg19 [35] using BWA-MEM algorithm [36]. Finally, we marked and removed the duplicates with Picard toolkit [37]. The observed variance being low, we did not apply the recalibration to the WES data.

We called the variants using Haplotype Caller (HC) algorithm from the GATK suite [31] in the targeted regions plus 150 bp up and downstream. Then, we filtered out SNVs and small indels (maximum length of 50 bp) according to the following criteria: total read depth DP < 12, mapping quality MQ < 30, variant confidence QD < 2 and strand bias FS score > 25. We annotated genotypes with an individual DP < 10 as missing before filtering out variants with call-rate < 95% with Plink1.9 [39]. Finally, we removed variants located in segmental duplications [40] and repeated regions, such as described in RepeatMasker from UCSC [41].

### Variant annotation and classification

We classified the remaining variants into rare variants (MAF < 1%), low frequency variants (1% ≤MAF<5%) and common variants (MAF ≥5%) according to their frequency in public databases. To evaluate these frequencies, we worked on 4 datasets extracted from: (1) the 1000 Genomes Project (2015 August); (2) the Exome Aggregation Consortium project (ExAC); (3) the Exome Sequencing project (ESP6500–6500 exomes); (4) the Complete Genomics project (CG69–69 individuals). We extracted the allele frequency in population with European ancestry when the information was available.

To investigate the predicted effect of the genetic variants on proteins, we annotated them with CADD phred-like score [42] using ANNOVAR [43] and with variant effect defined in SNPeff [44]. The former is a score obtained from a model trained to separate evolutionary conserved variants, likely deleterious, from simulated variants, likely benign. And the latter evaluate the putative variant impact at the transcript level by using sequence ontology.

### Principal Component Analysis of CEU controls and discovery set

To identify possible genetic population stratification between the discovery set and the 45 CEU controls, we performed a principal component analysis (PCA). For this analysis, we used a subset of variants located on autosomes classified as neutrals and frequents. We defined as neutral a variant annotated “LOW” by SNPEff and with a CADD phred-like score < 5. Those variants were considered frequent if their MAF was superior or equal to 5%. In addition, we processed the selected variants prior to performing the PCA analysis to remove those not in Hardy-Weinberg equilibrium (HW) and those in Linkage Disequilibrium (LD) with each other. We used Plink1.9 [39,45] to remove variants with HW_p-value_ < 0.01 and/or with LD ≥ 0.2. The Hardy-Weinberg equilibrium was evaluated on unaffected individuals only. We finally applied the PCA on these variations with the R package SNPRELATE [46].

### Selection of RA risk candidate variants

We first selected the rare variants (MAF ≤ 1%) observed in only one family with an in-house python script. Further, we chose variants segregating with RA within families and being absent from healthy relatives.

### Association analysis of selected genes with burden test

We tested the association of genes carrying at least one rare candidate variant using pVAAST [12]. This software extends the rare variant association test (RVAT) VAAST [47] to offer more power in the context of family-based studies. It provides a linkage-association score and a p-value for each evaluated gene and, a score for each variant carried by the gene to help the prioritization of these variants.

All the variants of our candidate genes, observed in the discovery set and/or in the 45 controls from 1000 genomes project, were included in the association analysis. Considering the heterozygous nature of our candidate SNVs, we tested the 73 candidate genes under a dominant model and set the maximum disease prevalence to 0.01, the world-wide prevalence of RA [48]. To evaluate the significance of the test, we authorized genotyping error and did not restrain penetrance value for gene-drop simulations. We estimated p-values by allowing pVAAST to perform up-to 10^6^ permutations.

### Validation in extended pedigrees of exome selected variants

The top 10 candidate variations after pVAAST analysis were re-sequenced in the validation set by using PGM^™^ System (Ion Personal Genome Machine^™^). Primers were designed with the Ion AmpliSeq^™^ Designer to target the selected variants. We mapped the reads to the reference genome with BWA [36] and recalibrated base score with BQSR tool from GATK suite [31]. We then called variants in the strict limits of the targeted regions with HC and applied the same quality filters used for WES to the observed variants. For samples in both discovery and validation sets, we checked the concordance with whole exome data using VCFtools program package [49].

### Genotyping of SUPT20H: *c.73T>A* in multiplex families and additional trios

The candidate rare variant was also genotyped using a custom assay with a FAM^™^ or VIC^®^ reporter Dye at the 5’end of each TaqMan^®^ MGB probe and a non-fluorescent quencher at the 3’end of each probe (Applied Biosystems, Foster City, CA, USA). Digital PCR (QX200™ Droplet Digital PCR, Bio-Rad Laboratories, California, USA) was used to detect variant alleles. First we analyzed members of the family in which the variant was characterized. Second, an independent sample of 188 trio families (one RA patient and his/her two parents from French European origin) was investigated to search for the variant, along with a positive and a negative control.

## Acknowledgments

We are grateful to RA patients and their families for participation in this study.

## Supporting information

**S1 Table:** Characteristics of 188 RA index cases in which the SNV c.73T>A carried by SUPT20H was investigated. Number of index cases / Number of index cases with data ^b^ previous and/or actual tobacco exposure (smokers and ex-smokers) RF: Rheumatoid Factor ACPA: Anti-Cyclic Citrullinated Peptide Antibodies

